# Structural determinants of inverted Alu-mediated backsplicing revealed by -MaP and -JuMP

**DOI:** 10.1101/2024.12.13.628372

**Authors:** Justin M. Waldern, Colin Taylor, Catherine A. Giannetti, Patrick S. Irving, Scott R. Allen, Mingyi Zhu, Rolf Backofen, Dave Mathews, Kevin M. Weeks, Alain Laederach

## Abstract

Biogenesis of circular RNA usually involves a backsplicing reaction where the downstream donor site is ligated to the upstream acceptor site by the spliceosome. For this reaction to occur, it is hypothesized that these sites must be in proximity. Inverted repeat sequences, such as Alu elements, in the upstream and downstream introns are predicted to base-pair and represent one mechanism for inducing proximity. Here, we investigate the pre-mRNA structure of the human *HIPK3* gene at exon 2, which forms a circular RNA via backsplicing. We leverage multiple chemical probing techniques, including the recently developed SHAPE- JuMP strategy, to characterize secondary and tertiary interactions in the pre-mRNA that govern backsplicing. Our data confirm that the antisense Alu elements, AluSz(-) and AluSq2(+) in the upstream and downstream introns, form a highly- paired interaction. Circularization requires formation of long-range Alu-mediated base pairs but does not require the full-length AluSq2(+). In addition to confirming long-range base pairs, our SHAPE-JuMP data identified multiple long-range interactions between non-pairing nucleotides. Genome-wide analysis of inverted repeats flanking circular RNAs confirm that their presence favors circularization, but the overall effect is modest. Together these results suggest that secondary structure considerations alone cannot fully explain backsplicing and additional interactions are key.

## Introduction

Circular RNAs (circRNAs) are formed by a noncanonical splicing process termed backsplicing (1–3). Backsplicing is a spliceosome-catalyzed process where a downstream splice donor (5’ splice site) is linked to an upstream splice acceptor (3’ splice site), covalently joining together one or more exons in a 5’ to 3’ phosphodiester linkage to form a complete circle (4, 5). CircRNA are often flanked by intronic inverted repeats, which have been proposed to drive backsplicing based on inter-intron RNA base-pairing via sequence complementarity (2, 3). In humans, 88% of circular RNAs are flanked by intronic Alu elements (6). Alu elements are non-autonomous retrotransposons that comprise approximately 10% of the human genome (7). Although Alu elements can be categorized into distinct familial lineages, the sequence similarity between any two Alu elements is typically greater than 80% (Stenger et al., 2001). The high degree of sequence similarity between Alu elements leads to a high degree of complementarity between inverted repeat Alu elements, which can base pair to form double-stranded RNA as evidenced by high degrees of ADAR editing in inverted repeat Alu elements in close genomic proximity (Bazak, Haviv, et al., 2014; Bazak, Levanon, et al., 2014). A leading hypothetical mechanism for backsplicing is that sense and antisense Alu elements, as inverted repeats, form inter-intronic structure across exons and drive backsplicing via sequence complementarity and long-range RNA structure that bring splice sites into proximity in three-dimensional space (3, 8).

The human homeodomain interacting protein kinase 3 (*HIPK3*) gene produces a single-exon circRNA from exon 2 that has been previously studied to decipher the sequence components required for backsplicing (2, 3, 9). The *HIPK3* exon 2 circRNA is flanked by intronic sense and antisense Alu elements, AluSz(-) and AluSq2(+), which are essential for backsplicing (Liang & Wilusz, 2014). However, regional deletions within these Alu elements can either ablate or retain backsplicing, suggesting that some portions of the Alu elements are essential for backsplicing whereas others are dispensable (2). Furthermore, when applying *in silico* energy minimization to predict Alu element structures, the thermodynamic stability of the truncated hairpin is not predictive of backsplicing, suggesting the possibility that multiple structural conformations or more complex structural interactions beyond simple base-pairing enable backsplicing (2). Distance requirements for backsplicing suggest there may be complex interactions at play; for example, moving the downstream AluSq2(+) up to 1,500 nucleotides away from the exon does not affect backsplicing, whereas moving the upstream AluSz(-) only 500 nucleotides away from the exon ablates backsplicing (10). Thus, based on sequence and predicted base-pairing considerations alone, it is difficult to predict if a given set of Alu elements will favor backsplicing. Furthermore, certain RNA binding proteins, such as ADAR and DHX9, specifically act on long double-stranded RNAs, such as paired inverted repeat Alu elements, and regulate backsplicing by targeting these structures (Aktaş et al., 2017). One aspect of pre-mRNA that remains understudied is its structure, and here we leverage novel chemical probing approaches to investigate the secondary and tertiary structures of the *HIPK3* pre-mRNA.

The size of human pre-mRNAs, where median intron lengths are over 1400 nucleotides (11), makes structural studies challenging. Chemical structure probing coupled to next-generation sequencing is one approach that can routinely probe structures of large RNAs (12, 13). However, most chemical structure probing techniques detect whether a nucleotide is paired (14, 15) but do not directly identify the pairing partner. SHAPE-JuMP (selective 2′-hydroxyl acylation analyzed by primer extension and juxtaposed merged pairs) leverages a highly processive reverse transcriptase coupled with a bivalent cross-linking reagent to identify paired and proximal nucleotides (16, 17). Here we leverage this strategy, coupled with conventional per-nucleotide chemical probing, to understand the structural determinants of backsplicing in the *HIPK3* circRNA. The integration of these data supports a complex structural model in which non-canonical base-pairing interactions in the pre-mRNA are as important in favoring backsplicing as inverted repeat base-pairing. Taken together, our data begin to reveal unexplored three-dimensional features of pre-mRNA structure and its role in regulating splicing.

## Results

### Mutational Profiling and Structural Analysis of inverted repeat pre-mRNA reveals Alu elements are likely structured

We begin our investigation using SHAPE-MaP (selective 2′-hydroxyl acylation analyzed by primer extension and mutational profiling) to chemically probe an *in vitro* construct of the human *HIPK3* pre-mRNA flanking exon 2. This construct is based on previous studies of the *HIPK3* circRNA, which has been described to circularize via backsplicing and form a 1099 nucleotide circle (2, 9).

When transcribed, the total construct is a 2926 nucleotide RNA that includes a 273 nucleotide antisense AluSz(-) in the upstream intron and a sense 301 nucleotide AluSq2(+) in the downstream intron (Figure 1A). Due to their inverted repeat configuration and high degree of complementarity, the two Alu elements are predicted to form base pairs across 85% of their sequence content.

**Figure 1:**
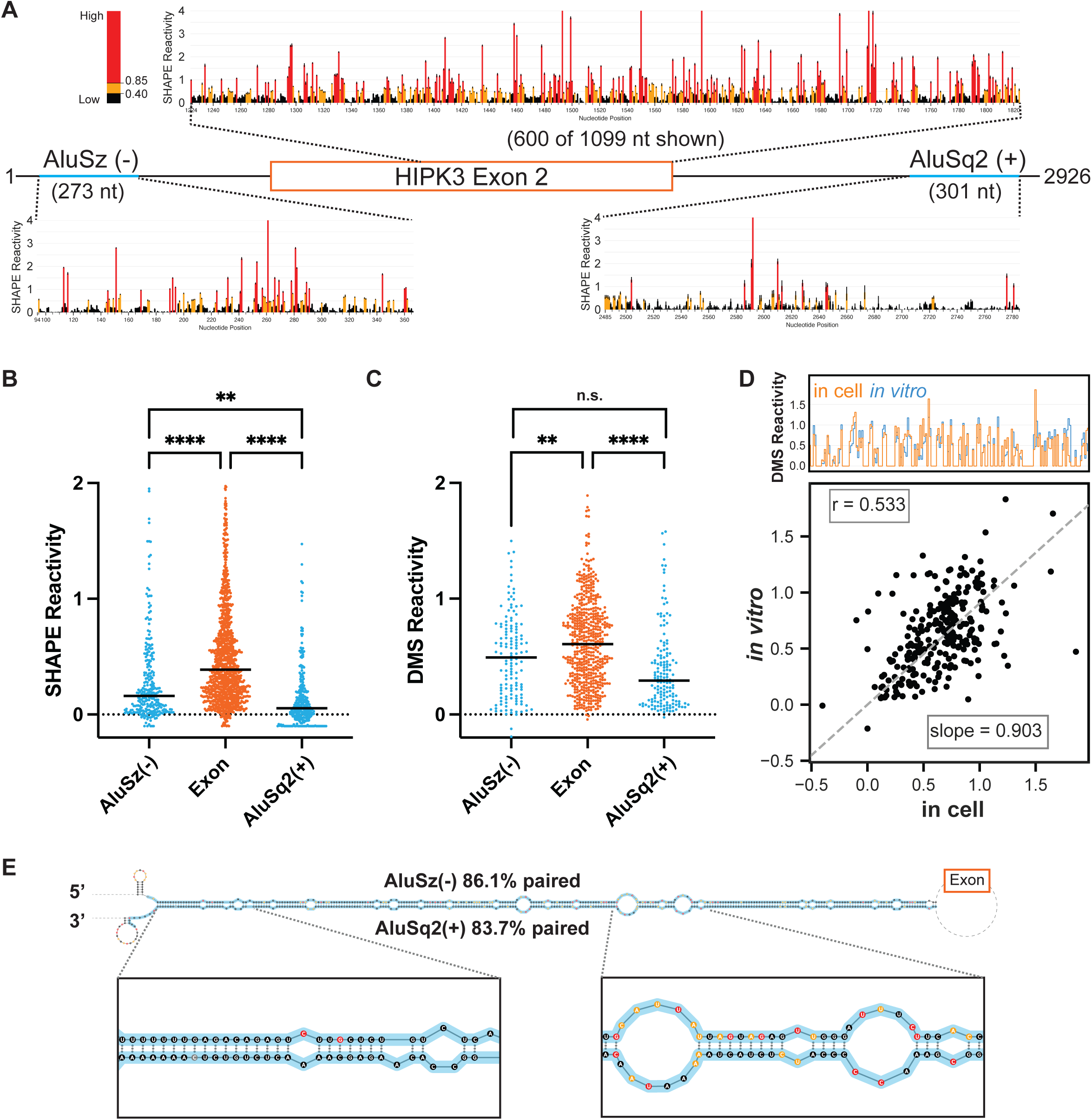
Chemical probing of *HIPK3* pre-mRNA. **A.** Pre-mRNA gene diagram and SHAPE profiles of *HIPK3* Exon 2 (orange) and its flanking Alu elements (blue). High SHAPE reactivities are shown in red, medium in orange, and low in black. **B.** Median regional normalized SHAPE reactivities comparing each Alu (blue) and the exon (orange). Significance determined with an ordinary one-way ANOVA. Significance throughout all figures: * p < 0.05, ** p < 0.01, *** p < 0.001, **** p < 0.0001 **C.** Median regional normalized DMS reactivities comparing each Alu (blue) and the exon (orange). DMS reactivity analysis is limited to adenosine and cytidine nucleotides. Significance determined with an ordinary one-way ANOVA. **D.** Comparison of normalized *in vitro* DMS reactivities with in-cell DMS reactivities. Top: skyline plot comparing per-nucleotide reactivity of a subset of the data. Bottom: Linear regression and correlation comparison of normalized reactivities. Correlation calculated as Pearson’s correlation coefficient. **E.** Maximum expected accuracy structure of the Alu-Alu hairpin informed by SHAPE probing. Structure is truncated for visualization and insets are zoomed in on representative regions. Alu elements are highlighted in blue and nucleotides are colored by SHAPE reactivity as in 1A. Percentage paired reflects the percentage of Alu nucleotides that are paired to any other nucleotide.

For SHAPE-MaP, we used the 5NIA (5-nitroisatoic anhydride) to probe the RNA and performed nucleotide-specific normalization (18) to compare median SHAPE reactivities across different regions of the pre-mRNA independent of nucleotide content. We performed two replicates of *in vitro* SHAPE-MaP experiments and analyzed the replicate correlation over several selected regions of the pre-mRNA (Supplementary Figure 1A) and found that the replicates are highly correlated (Pearson correlation coefficient r=0.83) (Supplementary Figure 1B and 1C). Overall, the SHAPE profiles of the *HIPK3* Alu elements show a consistently lower reactivity than that of the exon region, implying that the Alu elements are more structured than the exon (Figure 1A). Furthermore, the median SHAPE reactivities of both Alu elements are statistically significantly lower than that of the exon (Figure 1B). A similar trend is observed when considering the median *in vitro* adenosine and cytidine DMS (dimethyl sulfate) reactivities for the same construct (Figure 1C). The DMS data replicate with an r=0.79 (Supplementary Figures 1D and 1E).

In-cell pre-mRNA chemical structure probing data is more challenging to obtain, as introns are rapidly spliced and degraded. Using a tiled amplicon-based strategy, focusing on multiple amplicons targeting short regions, it is possible to collect chemical probing data at a limited scale (19). We designed and tested multiple amplicons spanning the *HIPK3* pre-mRNA and identified four amplicons from which we were able to obtain reproducible in-cell DMS probing data (r=0.7) (Supplementary Figure 1A, 1F, and 1G). Previous work has shown that the in-cell environment creates more biological noise (Smola et al., 2016), however the overall pattern of the two replicates here is the same (Supplementary Figure 1G). Comparing the in-cell and *in vitro* DMS probing data sets (Supplementary Figures 1H and particularly, 1I) reveals that reactivity in cells and *in vitro* are highly similar, suggesting that the *in vitro* data are representative of biologically relevant structures. Since the *in vitro* and in-cell probing data correlate well, we used the more comprehensive *in vitro* SHAPE-MaP data (Figure 1A) to model the structure of the full-length circularizing pre-mRNA. This model contains a unique structure of the AluSz(-)/AluSq2(+) paired interaction (Figure 1E), which is predicted to occur with high base-pairing probability (Supplementary Figure 2A). Similar results are obtained when predicting the structure using DMS data (Supplementary Figure

2B).

### Genomic and experimental measures of Alu-mediated long-range base-pairing and circRNA formation

Since it has been suggested previously that flanking inverted repeats favor circularization, we performed a genome-wide analysis of human circular RNAs and measured a higher log odds-likelihood of observing circularization if an exon is flanked by two inverted repeats compared to if an exon is flanked by a single or no Alu (Figure 2A). The difference in log-likelihoods is small, albeit statistically significant (p < 0.01). Although over 80% of circular exons are flanked by intronic Alu elements (6), there are Alu elements flanking an almost equally large proportion of non-circularizing exons. Thus, although circularization seems to be slightly more likely if flanking Alus are present, flanking Alu elements alone are not a strong predictor as to whether a circular RNA will form.

**Figure 2:**
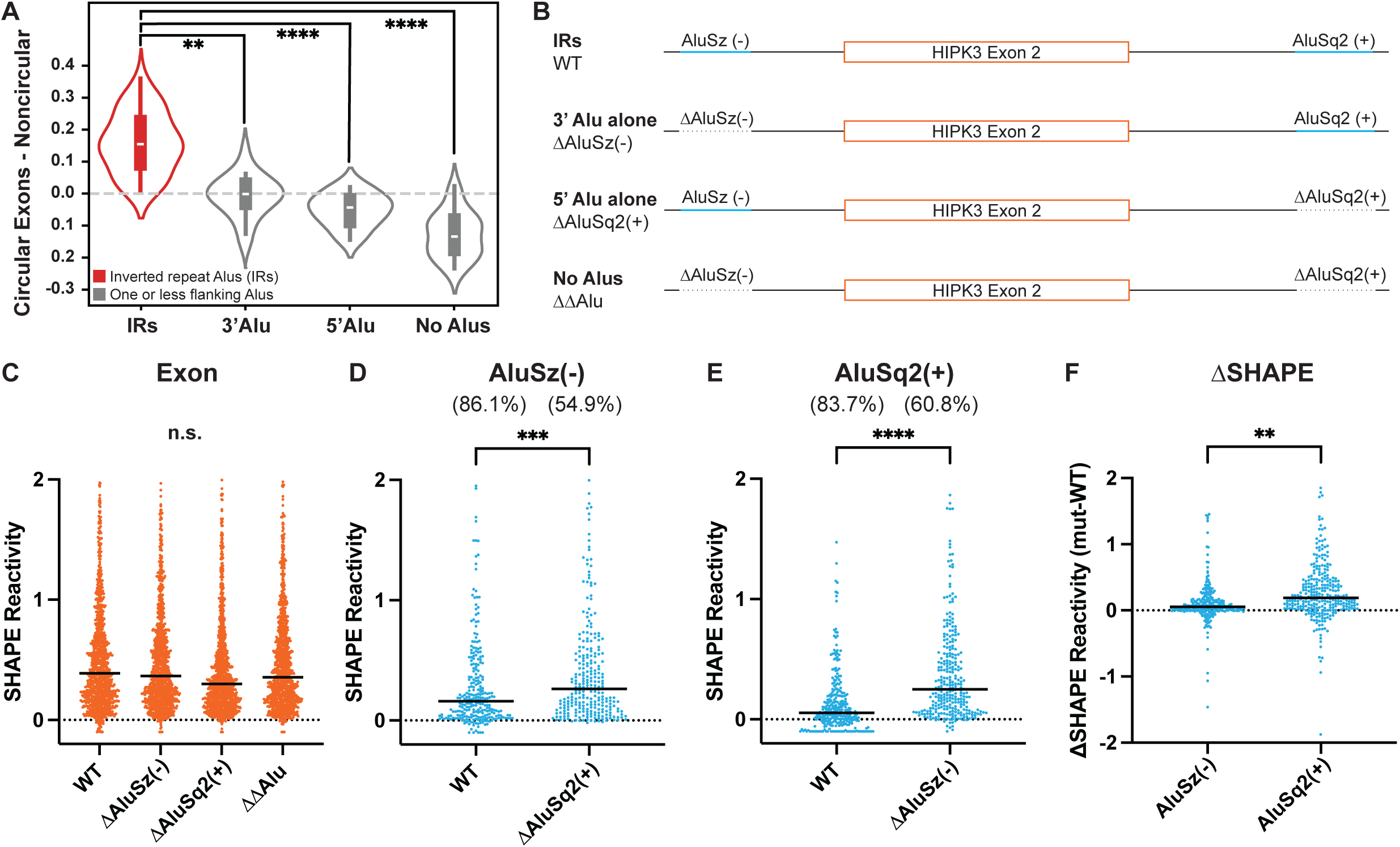
Inverted repeat Alu elements are required for circularization and likely base pair. **A.** Violin plot showing enrichment for inverted repeat Alu elements in exons found in the CircRNA database, CircBase (28), compared to exons not reported to circularize, grouped by the pattern of Alu elements flanking the exon. IR are inverted repeats (red), 3’ Alu lack a 5’ Alu upstream of the exon, 5’ Alu lack a 3’ Alu downstream of the exon, and No Alu have no Alu elements flanking the exon. Y-axis values are the difference in log odds ratios. Significance determined with Mann-Whitney U tests. Significance throughout all figures: * p < 0.05, ** p < 0.01, *** p < 0.001, **** p < 0.0001 **B.** Possible gene orientations for flanking Alu elements. Specifics for the *HIPK3* exon 2 system are shown. **C.** Median SHAPE reactivity of *HIPK3* exon 2 across multiple Alu contexts. Significance determined with an ordinary one-way ANOVA test. **D.** Median SHAPE reactivity of AluSz(-) with (WT) and without (ΔAluSq2(+)) its putative pairing partner. Numbers in parentheses are the percentages of AluSz(-) nucleotides paired in each context. Significance determined with a paired t-test. **E.** Median SHAPE reactivity of AluSq2(+) with (WT) and without (ΔAluSz(-)) its putative pairing partner. Numbers in parentheses are the percentages of AluSq2(+) nucleotides paired in each context. Significance determined with a paired t-test. **F.** Difference in SHAPE reactivities for each Alu, calculated by subtracting the SHAPE reactivity of each Alu in its putative pairing context (WT) from its SHAPE reactivity without its putative pairing partner (ΔAlu). Significance determined with an unpaired t-test.

Since the presence of flanking Alu elements does not guarantee backsplicing, we next focused on constructs that deleted one or both of the Alu elements flanking *HIPK3* exon 2 (Figure 2B) to experimentally investigate corresponding 3’, 5’ and No Alu, both structurally and functionally. We initially compared the median SHAPE reactivity for the 1099 nucleotide exon 2 for the WT and three constructs that deleted one or both of the flanking Alu elements (Figure 2C). Overall, the median SHAPE reactivity in the exon is not significantly different across all constructs (Figure 2C). In contrast, the median SHAPE reactivity of each Alu element is significantly higher if the opposite Alu is absent, as compared to the WT context (Figure 2D and 2E). The increase in reactivity without its pairing partner is more dramatic for AluSq2(+) than it is for AluSz(-) (Figure 2F). Similar trends are also observed in the DMS data (Supplementary Figure 3), suggesting each Alu is less paired when its partner is absent from the same pre-mRNA. Furthermore, when observed on a per nucleotide basis (Supplementary Figure 4) the difference in reactivity, as measured by ΔSHAPE (20, 21), is primarily driven by an overall change across the entire Alu element, although a few local regions do exhibit larger changes.

When using the SHAPE data to model the secondary structures of the ΔAluSq2(+) (Supplementary Figure 5) and ΔAluSz(-) (Supplementary Figure 6) deletion constructs, inter-intron pairs are no longer predicted as expected. Instead, the remaining Alu element in each deletion construct is predicted to form interactions between the Alu element and its flanking intronic sequences (Supplementary Figures 5 and 6A-C). Folding the AluSq2(+) sequence in isolation combined with the ΔAluSz(-) SHAPE data provides a more traditional Alu-like fold (Supplementary Figures 6D and E). Overall, the structural data and reactivity changes in the deletion mutants indirectly suggest long-range base-pairing interactions are occurring in our WT pre-mRNA construct and that deletion of either Alu (or both) eliminates inter-intron base-pairing.

### Functional characterization of all constructs and partial deletions of AluSq2(+)

To examine the effects of the potential pairing interaction on backsplicing, we designed a qRT-PCR assay to measure the circularization efficiency of different constructs (Supplementary Figure 7). None of the deletion mutants – ΔAluSz(-), ΔAluSq2(+) or ΔΔAlu – produce circles (Figure 3A), emphasizing that circularization is dependent on the presence of the both AluSz(-) and AluSq2(+), consistent with previous studies (2). To identify the length of the Alu-Alu base-pairing interaction required for circularization, we designed partial deletions of AluSq2(+), where 157 or 189 nucleotides are deleted from the 5’ end of AluSq2(+). The partial deletion constructs, Δ157AluSq2(+) and Δ189AluSq2(+), circularize with similar efficiency as WT (Figure 3A). These results suggest that circularization is dependent on inter-intron base-pairing but that partial length inverted repeats are sufficient. To further identify if circularization requires a specific length of Alu-Alu pairing interaction, we performed a human genome-wide analysis of circularizing and non-circularizing exons flanked by inverted Alu repeats and plotted the distribution of the length of the downstream Alu (Figure 3B). We observed no difference in the distributions, consistent with our HIPK3 circularization assay and our partial AluSq2(+) deletion mutants. Together, these results suggest that approximately 100 nucleotides of complementary sequence are sufficient to favor circularization, and that two full-length Alu repeats are not essential.

**Figure 3:**
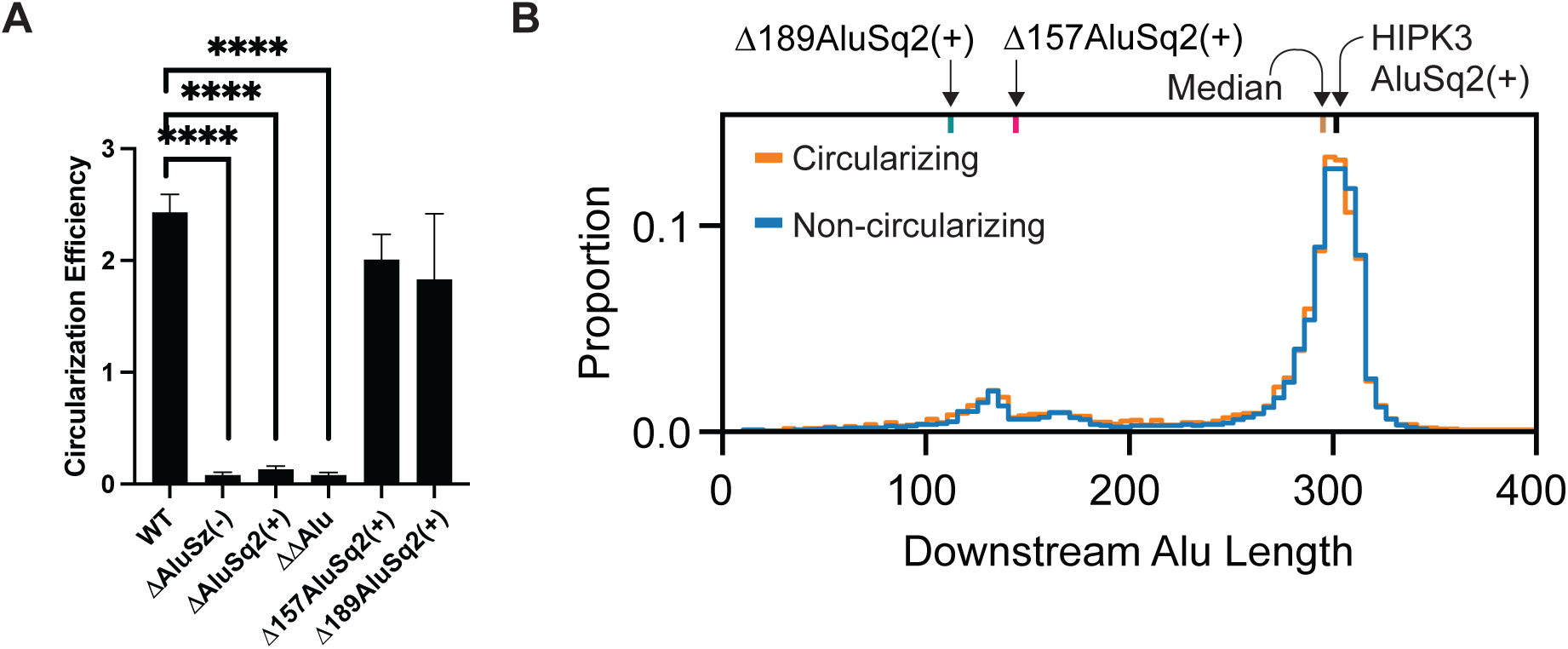
Alu elements are required and partial Alu elements are sufficient for circularization. **A.** Circularization efficiency of *HIPK3* exon 2 in differing Alu contexts. Significance determined with an ordinary one-way ANOVA and Dunnett’s multiple comparisons test. Significance throughout all figures: * p < 0.05, ** p < 0.01, *** p < 0.001, **** p < 0.0001 **B.** Distribution of the length of Alu elements downstream of exons. Alu elements associated with exons in the CircRNA database, circBase, are shown in orange, whereas Alu elements associated with exons not known to circularize are in blue.

### Direct probing of long-range interactions using SHAPE-JuMP

The structural data here are consistent with a model in which the HIPK3 exon 2 flanking AluSz(-) and AluSq2(+) form an extended secondary structure. To directly measure the Alu-Alu interaction, we used SHAPE-JuMP (-Juxtaposed Merged Pairs) (16, 17) to obtain nucleotide resolution information on long-range interactions in the WT, Δ157AluSq2(+), Δ189AluSq2(+), ΔAluSz(-) and ΔAluSq2(+) constructs. In SHAPE-JuMP experiments, a chemical cross-linker, (here, a psoralen crosslinker) covalently links nucleotides that are in close three-dimensional proximity. The crosslinks are then “read” using a highly processive reverse transcriptase that “jumps” across the crosslinks, recording the crosslinked interaction as a deletion. The resulting cDNA is sequenced and deletion rates for specific nucleotide pairs are counted by massively parallel sequencing (16, 17). We used an amplicon-based strategy to capture long-range interactions using short read sequencing and to specifically capture inter-intron jumps.

We first evaluated the cumulative ranked distribution of observed reverse transcriptase jump frequency (deletions normalized to read depth) for the five probed constructs (Figure 4A). The highest jump frequency is observed for the WT and Δ157AluSq2(+). The Δ189AluSq2(+) construct has an intermediate level of jumps, whereas both ΔAluSq2(+) and ΔAluSz(-) have very few jumps that are essentially indicative of background noise. The jump frequency for each construct is consistent with its circularization efficiency, suggesting that the structures detected by -JuMP are important for circularization. We performed -JuMP experiments with circularizing constructs in replicate (Supplemental Figure 8); in subsequent analyses, both replicates have been combined into a single data set. We use a “triangular” representation to visualize the -JuMP data in the context of the secondary structure model (Figure 4B) (22). Several important features of the SHAPE and DMS informed structural models for the WT construct are clear when superimposed over each other and colored by pairing probability (Figure 4B). Most of the structures observed (dots representing base pairs) are short-range local structures, as indicated by their positions near the top of the graph. Among these small local structures, there is some variance between structures that are SHAPE-supported, DMS-supported, or both. However almost all the dots representing the AluSz(-)/AluSq2(+) interaction are black (near the bottom point of the triangle diagram), confirming that the SHAPE and DMS informed base-pairing probabilities are both in agreement and very high (approximately 1.0) (Figure 4B, enlarged in Figure 5). We observe no alternative base-paring in either the SHAPE or DMS informed structures in proximity to the AluSz(-)/AluSq2(+) pairs (Figure 4B). Importantly, there are no long-range base-pairing interactions connecting the splice sites or near where the ends of the exon would come together for backsplicing (the tip of the orange triangle), suggesting that base-pairing near the backsplice junction is not occurring; instead, the only predicted long-range base-pairing is the AluSz(-)/AluSq2(+) interaction.

**Figure 4:**
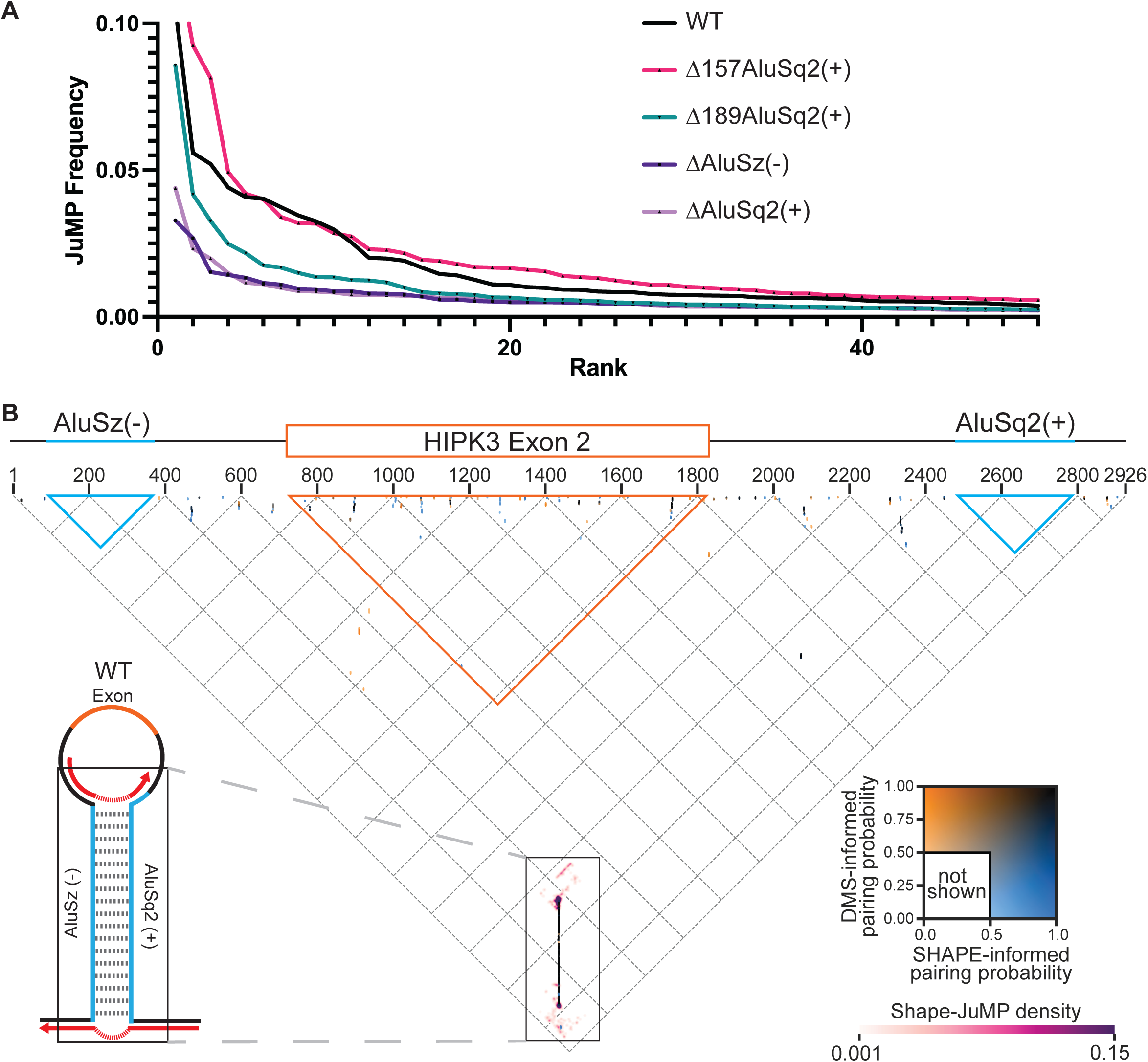
JuMP, SHAPE, and DMS data are in agreement. **A.** JuMP event frequency distribution, where the JuMP frequency is plotted on the y-axis sorted from highest to lowest frequency (rank, x-axis). The amount of high frequency JuMPs corresponds to the amount of predicted Alu pairing. **B.** The inverted triangle represents all possible pairings for the WT *HIPK3* construct, where the horizontal axis is the 5’ to 3’ sequence, as illustrated by the gene diagram (top). The blue triangles represent the space occupied by each Alu element, whereas the orange represents the exon. Dots on the triangle represent predicted base-pairing interactions, colored by pairing probability informed by each probe (DMS or SHAPE) as seen in the inset color key on the right. Interactions based on JuMP data are represented by heatmaps (pink clouds) and shown as JuMP density.

**Figure 5:**
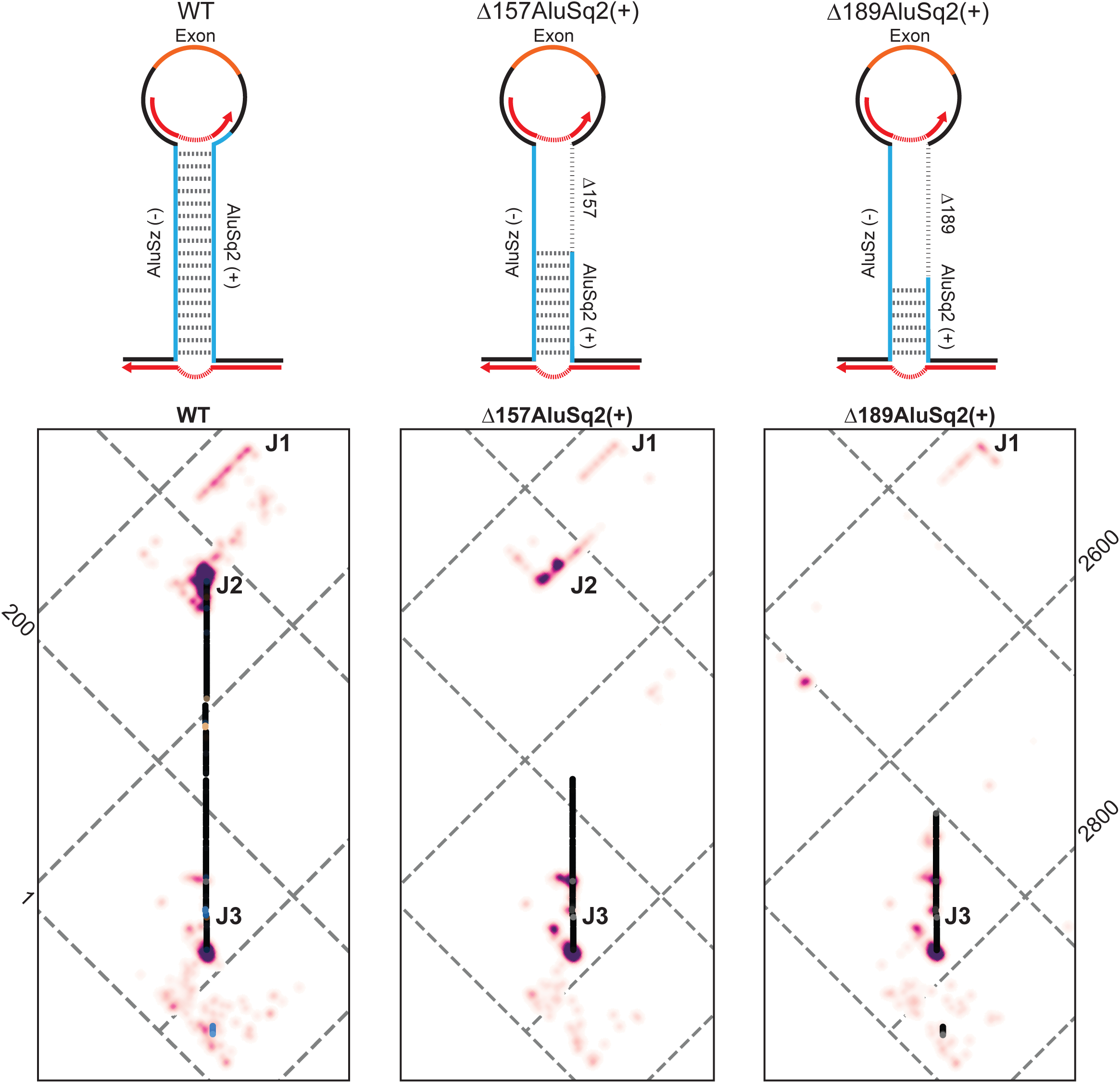
JuMP data of circularizing constructs mapped to WT coordinates show broad structural agreement. Top, model structure illustrations for wild type (WT) and partial Alu deletions, Δ157AluSq2(+) and Δ189AluSq2(+). Alu elements are illustrated in blue with deleted sequence marked by a dashed line and the exon in orange. Red arrows represent amplicon-targeted predicted JuMP events, with the deletion marked by the dashed portion. **B.** Zooming in on the Alu-Alu interaction of wild-type (WT), Δ157AluSq2(+), and Δ189 AluSq2(+) structures, mapped to WT coordinates. JuMP density is colored as in Figure 4B, as is pairing probability for WT. Pairing probability for Δ157AluSq2(+) and Δ189AluSq2(+) are greyscale colored based only on SHAPE-informed pairing probabilities. Notable JuMP interactions are denoted by J1, J2, and J3.

Therefore, the -JuMP experiments were designed to target the predicted AluSz(-)/AluSq2(+) interaction in the WT construct. The most prominent jumps are observed at both ends of the AluSz(-)/AluSq2(+) hairpin (labeled as J2 and J3) in the WT construct (Figures 4B and 5). These experiments exhibit very few inter-Alu crosslink jumps inside the hairpin; most jumps are concentrated in non-paired regions (J1) and at the very ends of the paired region (J2 and J3) (Figure 4B). These observations are consistent with the ability of SHAPE-JuMP to detect both secondary structures and tertiary (non-base-pairing) interactions (16, 17). The detection of jumps at both ends of the AluSz(-)/AluSq2(+) hairpin provide direct evidence for the existence of this long-range interaction.

When comparing the three constructs that circularize (WT, Δ157AluSq2(+), and Δ189AluSq2(+)) by mapping the deletion mutants to WT coordinates, there is a consistent pattern of jumps that directly detect and further support the existence of the Alu-Alu hairpin (Figure 5). All three structures show nearly identical JuMP interactions that directly indicate Alu-Alu pairing (J3) (Figure 5). The construct with the shortest Alu-Alu interaction (Δ189AluSq2(+)) shows the most internal Alu-Alu hairpin jumps (Figures 5 and 6), suggesting that the lack of inter-Alu jumps deep in the Alu-Alu hairpin in the WT construct is likely due to the extremely high level of crosslinking and significant base-pairing of two full length inverted Alu elements, as these factors limit the processivity of the reverse transcriptase resulting in fewer sequencing reads deep into the hairpin.

**Figure 6:**
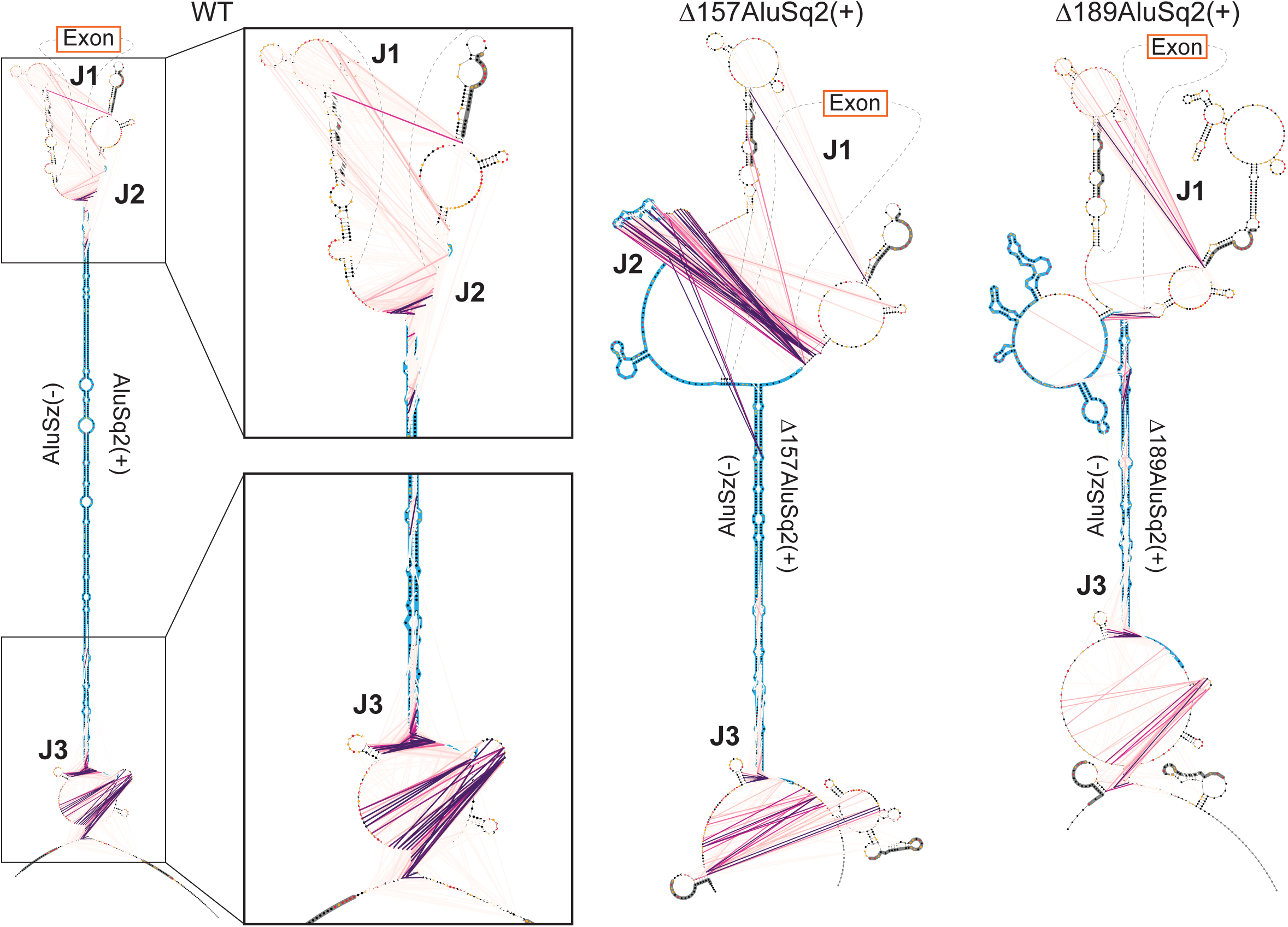
Maximum expected accuracy secondary structure diagrams of the Alu-Alu interaction of wild-type (WT), Δ157AluSq2(+), and Δ189AluSq2(+) constructs. Structures are informed by SHAPE chemical probing. Alu elements are highlighted in blue, with primer locations from JuMP highlighted in dark gray. JuMP interactions are shown by pink/purple lines, with the color scale from Figure 4B. Individual nucleotides are colored by SHAPE reactivity. Structures are truncated to focus on Alu elements and JuMP interactions. Notable JuMP interactions are denoted by J1, J2, and J3 and correspond to the same labels in Figure 5.

The patterns of jump density for the WT, Δ157AluSq2(+), and Δ189AluSq2(+) are broadly similar for non-inter-Alu pairing (Figure 5). Specifically, there is a very similar pattern of density at the bottom of the AluSz(-)/AluSq2(+) for all three constructs (J3) extending beyond the hairpin and representing the areas nearest the 5’ and 3’ ends of the constructs (Figure 5). All three also show a similar pattern of jumps on the exon-proximal side of the Alu elements (J1) (Figure 5). These regions lack sequence complementarity and are not predicted to be paired in any of the structure models, suggesting the crosslink jumps are reporting three-dimensional spatial proximity in these regions.

The same data can also be visualized on more traditional secondary structure diagram models of each circularizing construct (WT, Δ157AluSq2(+), and Δ189AluSq2(+)) (Figure 6). These secondary structure diagrams reveal the extent to which many of the observed jumps are not between canonically base-paired regions. Interestingly, despite the removal of 157 nucleotides in AluSq2(+), the J2 interaction at the exon-proximal end of the Alu elements is maintained across WT and Δ157AluSq2(+). The removal of an additional 32 nucleotides in Δ189AluSq2(+) abolishes this J2 interaction, however -JuMP still detects the base pairing at both ends of the truncated AluSq2(+) in Δ189AluSq2(+) (Figure 6). In effect, the AluSz(-)/AluSq2(+) hairpin brings into close three-dimensional proximity two intronic regions (J1 and J3) causing them to interact spatially. These interactions are preserved across all three circularizing constructs (WT, Δ157AluSq2(+), and Δ189AluSq2(+)). Thus, our data suggest that the ability of inverted repeats to lock two introns together is spatially orienting the 5’ and 3’ splice sites which facilitates backsplicing.

## Discussion

Inverted repeats like the AluSz(-)/AluSq2(+) interaction in *HIPK3* exon 2 appear to form highly stable, fully paired interactions, even though they are over 2000 nucleotides apart. These long-range interactions are directly supported by extensive chemical probing data in our study. These extended pairing interactions, in turn, create sufficient proximity between intronic structured elements that they are detectable by chemical crosslinking and SHAPE-JuMP. This spatial proximity is sufficient to facilitate backsplicing. This work represents the first direct experimental detection and structural description of a long-range Alu-Alu interaction at single-nucleotide resolution.

In this work, we experimentally explored the pre-mRNA structures that facilitate backsplicing in the *HIPK3* exon 2 circular RNA. We coupled this experimental work with genome-wide analyses that further support a complex, and likely structural, explanation of backsplicing beyond inverted sequence complementarity. Genomic analyses suggested that the presence of exon flanking inverted repeat Alu elements is only modestly predictive of exon circularization, as the log-likelihood of circularization is only marginally larger for exons flanked by inverted repeats versus not (Figure 2A). Furthermore, the length of the predicted Alu-Alu interaction is insufficient to identify those exons capable of circularizing (Figure 3B). Combined with the structural data presented here, the presence and length of inverted repeat Alu elements are not strong predictive features of circularization alone. Our experimental data reveals a more complex induced proximity that is conserved in all circularizing constructs.

The structures of inverted repeats present a unique challenge in structure prediction. Inverted repeat Alu elements are highly complementary and are predicted by traditional thermodynamic modeling to form large hairpins even when separated by long stretches of sequence. In the wild-type *HIPK3* construct, the AluSz(-)/AluSq2(+) interaction is predicted in the minimum free energy structure and also has 100% base-pairing probability in the Boltzmann suboptimal ensemble. The high degree of complementarity drives enormous thermodynamic favorability that excludes the possibility of alternative structures, despite the large distance between the inverted repeats in traditional thermodynamic models.

It has long been understood that Alu elements comprise approximately 10% of the human genome (7); nonetheless, the abundance of repetitive sequences in the human genome has only recently been characterized fully, as part of the Telomere to Telomere (T2T) genome (23). Given that a majority of the genome is also transcribed, the potential for inverted repeat base-pairing in the transcriptome is enormous. These inverted repeat interactions have the potential to bring diverse RNA sequences, nominally kilobases apart in primary sequence, into physical proximity. Determining the specificity with which two inverted repeats form a long-range interaction is challenging for repetitive sequences. In our experiments, we leveraged the unique ability of the -JuMP RT, coupled with specific priming outside of the repetitive sequences, to directly measure this interaction. Our experiments also revealed persistent non-base-pairing interactions outside the complementary regions. These data suggest that non-canonical through-space interactions in a pre-mRNA may be common and imply that secondary structure considerations alone are insufficient to understand and ultimately predict specific aspects of splicing regulation.

## Materials and Methods

### Construct design

HIPK3 constructs were initially designed based on (2). Constructs were synthesized by Genscript to insert the HIPK3 exon 2 and its surrounding sequence (GRCh38/hg38 chr11:33,285,723-33,288,525) into a pcDNA 3.1 backbone. Mutant derivatives of this construct (ΔAluSz(-), ΔAluSq2(+), ΔΔAlu, Δ157AluSq2(+), and Δ189AluSq2(+)) were also generated by Genscript and full sequence files can be found in Supplementary Files 1-6.

### Cell culture and Transfection

HeLa cells (ATCC number CCL 2) were cultured in high glucose DMEM (Gibco) supplemented with10% fetal bovine serum (Sigma) and 0.5 % pen/strep (Gibco). Cells were maintained at 37° C and 5% CO2.

Transfections were performed with Lipofectamine 3000 (Invitrogen) per manufacturer protocols. Cells were seeded 24 hours prior to transfection, where1 µg of plasmid was transfected per well in 6-well plates. Media was exchanged for fresh media 24 hours post transfection, and cells were harvested 48 hours post transfection.

### RNA preparation (*in vitro*)

DNA template for *in vitro* transcription was generated with PCR (primers JW25 and JW15 in Supplementary File 7) using the construct plasmids as a template and Q5 Hot Start High-Fidelity DNA polymerase (NEB). PCR reactions were cleaned up with the Monarch PCR & DNA Cleanup kit (NEB), with product size and integrity verified by agarose gel electrophoresis. RNA was *in vitro* transcribed from the clean PCR product using HiScribe T7 High Yield RNA Synthesis kit (NEB), followed by TurboDNase treatment (Invitrogen). RNA was isolated by ethanol precipitation.

### Chemical probing - *in vitro* SHAPE

Selective 2’hydroxyl acetylation analyzed by primer extension and mutational profiling (SHAPE-MaP) with 5-nitroisatoic-anhydride (5NIA) was used to probe RNA structure at all four nucleotides. To begin, 3 µg of *in vitro* transcribed (IVT) RNA was diluted in a total volume of 50 µL and denatured (65° C for 5 minutes, followed by ice for 2 minutes). Denatured RNA was combined with 50 µL 2X Bicine folding buffer (600 mM Bicine pH8.3, 300 mM NaCl, 10 mM MgCl2) and 1 µL RNase inhibitor, then refolded at 37° C for 10 minutes. RNA was then added to 10 µL of 250 mM 5NIA (in DMSO) or 10 µL DMSO as a control and incubated at 37° C for an additional 10 minutes, until finally quenching on ice. RNA cleanup was performed with RNAClean XP beads (Beckman Coulter).

RNA was reverse transcribed under Mutational Profiling Reverse Transcription conditions (MaP-RT) with Superscript II (Invitrogen), randomly primed with random nonamers. Briefly, 30 µL of RNA was premixed with 1 µL of 200 ng/µL Random Primer 9 (NEB) and 2 µL 100 mM dNTPs (NEB) and heated at 65° C for 5 minutes, then chilled on ice for 2 minutes. Added to each reaction was 4 µL of first strand buffer (0.5 M Tris pH 8.0, 0.75 M KCl), 4 µL of 100 mM DTT, 0.48 µL 100 mM MnCl2, 0.5 µL RNase inhibitor (NEB), and 2 µL Superscript II. Reactions were carried out with the following program: 25° C for 2 minutes, 42° C for 3 hours and 70° C for 15 minutes. Reverse transcription reactions were cleaned up with G50 columns (Cytiva).

Second strand synthesis was performed with the NEBNext Ultra II Non-Directional RNA Second Strand Synthesis Module (NEB). Libraries were generated with the NEBNext Ultra II FS DNA Library Prep Kit for Illumina (NEB). Library concentration and purity was verified using the Qubit dsDNA HS kit (Invitrogen) and the Bioanalyzer 2100 (Agilent), followed by equimolar pooling and sequencing on the Illumina MiSeq platform.

### Chemical probing - *in vitro* DMS

For structure probing with dimethyl sulfate (DMS), 3 µg of IVT RNA was diluted in 15 µL of molecular biology grade water and heat denatured (65° C for 5 minutes, followed by ice for 2 minutes). Denatured RNA was combined with 15 µL 2x Bicine folding buffer and refolded at 37° C for 30 minutes. 9 µL of folded RNA was then added to either 1 µL of 1.7 M DMS (final concentration 170 mM) prepared in a 1:2 (v/v) nitromethane/sulfolane (NS) solution or to 1 µL of NS solution as a vehicle control as in (24). The reaction was incubated at 37° C for 6 minutes before quenching with an equal volume of 1:3 (v/v) beta-mercaptoethanol (BME) in water. Reactions were buffer exchanged with G50 spin columns (Cytiva) prior to MaP-RT.

DMS-optimized MaP-RT was performed as previously described (21). Briefly, 8.8 µL of the probed RNA were combined with 2 µL 10 mM dNTPs (NEB) and 1 µL 200 ng/µL random nonamer (Random Primer 9, NEB). This mixture was heat denatured at 98° C for 1 minute, then chilled on ice for 2 minutes. Added to each reaction was 2 µL of freshly prepared 10x NTP minus first strand buffer (0.5 M Tris pH 8.0, 0.75 M KCl, 0.1 M DTT), 4 µL 5 M betaine, and 1.2 µL 100 mM MnCl2. Reactions were incubated at 25° C for 2 minutes before adding 1 µL of either SuperScript II RT enzyme or water (for no RT controls). Reactions were carried out with the following program: 25° C for 10 minutes, 42° C for 90 minutes, 10 cycles of 50° C for 2 minutes followed by 42° C for 2 minutes, then 70° C for 10 minutes and a hold at 12° C. Reactions were cleaned with G50 spin columns (Cytvia) prior to second strand synthesis. Second strand synthesis and library preparation was performed as described above for the 5NIA treatment.

### Chemical probing - in cell DMS

HeLa cells were transfected in 6-well plates with 1 µg of wild-type HIPK3 plasmid, as described above. At harvesting (48 hours post transfection), cells were washed with 1X phosphate buffered saline (PBS, Gibco) and allowed to equilibrate at 37° C in buffered media (DMEM, 200 mM Bicine pH 8). Each well was treated with 100 µL of either 100% ethanol as a control, or DMS diluted 1:20 in 100% ethanol at 37° C for 6 minutes. To quench the reaction, 1 mL of cold 20% BME was added to each well. After removing all liquid waste, RNA was isolated from cells with Trizol Reagent (Invitrogen) following the manufacturer’s protocol. To remove DNA, RNA was treated with TurboDNase (Invitrogen) following manufacturer’s protocol, with the modification of treating for 1 hour, adding 1 µL additional DNase halfway through. Following DNase digestion, RNA was purified by ethanol precipitation. For MaP-RT, 2 µg of RNA was used as template and reverse transcription was performed as described above for *in vitro* DMS probing. RT reactions were cleaned up with G50 columns (Cytivia) Library preparation for in cell probing followed an amplicon-based strategy with 4 different primer sets (Supplementary Figure 1A and Supplementary File 7). PCR-1 was performed with Q5 Hot Start High-Fidelity DNA polymerase (NEB) using the entire 20 µL MaP-RT reaction and following manufacturer’s protocol with 20 cycles of amplification. PCR-1 product was cleaned up with Ampure XP bead-based reagent (Beckman Coulter) following manufacturer’s protocol. Product concentration was evaluated with Qubit dsDNA HS kit (Invitrogen) to determine input for indexing PCR (PCR-2) and rule out DNA contamination in no-RT controls. For PCR-2, 25 ng of input DNA were amplified using NEBNext Multiplex Oligos for Illumina (NEB) and NEBNext Ultra II Q5 Master Mix (NEB) according to manufacturer’s protocol with half as much primer (0.25 µM final concentration) and 10 cycles of amplification. Products were cleaned up as in PCR-1, and purity verified with the Agilent 4200 TapeStation System. Libraries were pooled at equimolar concentrations and sequenced with the Illumina NextSeq 1000 platform.

### Structure probing data analyses

Structure probing data was primarily analyzed using Shapemapper2 v2.1.5 (25) and RNAvigate v1.0.0 (26), with structure prediction from RNAstructure Version 6.4 (27). For 5NIA data, the SHAPE profiles were rescaled according to (18) and values less than -0.1 were set to -0.1 since any values less than zero have fewer mutations in the treated than the untreated control and are functionally zero. For DMS analyses, the input fasta file was modified to mask guanosine and uridine into lowercase and only analyze adenosine and cytidine nucleotides. SHAPE and DMS profiles, skyline plots, and correlation plots were generated with RNAvigate. Correlation coefficients were calculated with RNavigate. Minimum free energy structures were generated with Fold from the RNAstructure package. Pairing probabilities were generated with the partition function in RNAstructure followed by the ProbabilityPlot function. Maximum expected accuracy structures were generated using the MaxExpect function in RNAstructure. Structures were arranged for visibility and organization in StructureEditor 1.0, then plotted with associated data and annotations in RNAvigate.

### Genome-wide analysis

Circular RNA coordinates were collected from the circBase database (28). Coordinates of all exons were collected from GENCODE v44. Alu elements coordinates were identified from RepeatMasker annotations in Hg38 (29). To categorize exons into circularizing and non-circularizing, annotations from circBase containing exactly 1 exon were intersected using BedTools Intersect (30) with annotations from GENCODE v44, with exons containing an overlap marked as circularizing.

Exon-flanking Alus were defined as full-length Alu elements within 2000 nucleotides of an exon. Therefore, exons were considered to have a 5’ flanking Alu if there was an Alu element that met the definition criteria on the 5’ side but not the 3’ side. Exons were considered to have a 3’ flanking Alu if there was an Alu element that met the definition criteria on the 3’ side but not the 5’ side. Exons were considered to have no Alu elements if no Alus meeting the definition criteria were found on either side. Exon-flanking Alus were considered inverted repeats if two Alus met the criteria on either side of the exon, and one Alu was found on the positive strand with the other Alu on the negative strand. Differences between groups were evaluated by taking the difference in the log odds ratio and compared statistically with Mann-Whitney U tests. To produce Alu element length distributions, downstream Alus were categorized as described above, regardless of length, into circularizing and non-circularizing and plotted based on length.

### Circularization Assays

For circularization assays, constructs were transfected into 6-well plates as described above. RNA was isolated using the Quick RNA Miniprep kit (Zymo) with on column DNase treatment, followed by an additional DNase treatment with TurboDNase (Invitrogen), and cleaned up again by ethanol precipitation. Preliminary circularization assays were performed with RT-PCR and evaluated using gel electrophoresis.

RNA was subjected to qRT-PCR analysis with custom Taqman gene expression assays (Life Technologies). Reverse transcription was performed with SuperScript IV VILO Master Mix (Invitrogen) with ezDNase treatment using 500 ng RNA. Three different custom Taqman probes (illustrated in Supplementary Figure 7) were designed with the LifeTechnologies custom gene expression assay design tool corresponding to circular RNA (assay ID: APFVV29), pre-mRNA (assay ID: APGZPM6), and lastly the Neomycin resistance gene (assay ID: APEP2NU), which is specific to the plasmid and absent from the mammalian genome. The qPCR reactions were performed using the TaqMan Fast Advanced Master Mix for qPCR (Applied Biosystems) in 10 µL reactions (4.5 µL diluted cDNA, 5 µL 2X Master Mix, and 0.5 µL TaqMan probe). For each qPCR reaction, 2 ng of cDNA per well was used for the circular and pre-mRNA probes, whereas 0.2 ng of cDNA per well was used for the Neomycin probe. qPCR was performed on the Applied Biosystems QuantStudio 6 Flex Real-Time PCR System.

For data analysis, the Neomycin resistance gene was used as a reference gene for qPCR to account for any variation in transfections. To quantify normalized circular and pre-mRNA, relative expression was calculated as 2^-ΔCt^, where ΔCt is the difference between the target and the reference (Neomycin resistance). Circularization efficiency was calculated by taking the ratio of normalized circRNA to pre-mRNA and standard deviations were calculated with propagated error.

### SHAPE-JuMP probing

For SHAPE-JuMP probing,1 µg of *in vitro* transcribed RNA was denatured (65° C for 5 minutes and snap cooled on ice, 2 minutes) before being refolded in 2X Bicine folding buffer at 37° C for 15 minutes, followed by addition of 0.25 µg 4’- aminomethyltrioxsalen hydrochloride (AMT) per reaction (water for untreated) and incubation at 37° C for an additional 15 minutes. Samples were then crosslinked on ice with UV light at 365 nm for 10 minutes. Reactions were cleaned up with G50 columns (Cytiva).

For reverse transcription, 1 µL RT primer (JW39 or JW41, Supplementary File 7) was mixed with 9 µL of crosslinked RNA, denatured at 65° C for 5 minutes, followed by snap cooling on ice for 2 minutes. The RNA/primer mix was added to a reverse transcription reaction with 10 µL 5X ThermoPol buffer made fresh (100 mM Tris-HCl pH8.8, 50 mM (NH4)2SO4, 50 mM KCl, 0.5% Triton X-100, 10 mM MgCl2), 1 µL 10 mM dNTPs (NEB), 28 µL molecular biology grade water, 1 µL C8 enzyme. Reverse transcription was performed at 65° C for 4 hours as previously described (17). Reverse transcription reactions were cleaned up with G50 columns (Cytiva).

Library preparation for JuMP experiments followed an amplicon-based strategy. PCR-1 (JW45&JW49 or JW48&JW50, Supplementary File 7) was performed with Q5 Hot Start High-Fidelity DNA polymerase (NEB) following manufacturer’s protocol for 25 cycles of PCR. PCR-1 reactions were cleaned up with Ampure XP bead-based reagent (Beckman Coulter) and concentrations were checked with Qubit dsDNA HS kit (Invitrogen). For PCR-2, 1 ng of PCR-1 product was used as template with TruSeq indexing primers (Illumina) and 10 cycles of PCR. PCR-2 product was cleaned up with Ampure XP bead-based reagent (Beckman Coulter). Library concentration and purity was verified using the Qubit dsDNA HS kit (Invitrogen) and the Bioanalyzer 2100 (Agilent), followed by equimolar pooling and sequencing on the Illumina MiSeq platform.

Sequencing data from SHAPE-JuMP experiments were analyzed with ShapeJumper V1.0 (16). SHAPE-JuMP frequencies were calculated with using the data from normalizeDeletionRates.py in the ShapeJumper package, with low count (<20) JuMP events omitted. Low count JuMP events were omitted because they can be artificially high in this system due to both a low JuMP count and a low read depth (i.e., 1 event in 1 read is a 100% frequency), where the low read depth occurs on JuMPs deep into the read from the poor processivity of the reverse transcriptase. Values plotted are the differences in frequencies between the treated and untreated samples.

## Acknowledgments

We wish to thank William F. Marzluff for useful discussions. This work was supported by U.S. National Institutes of Health grants R01 HL111527 (to A.L. and K.M.W.), R35 GM140844 (to A.L.), and R35 GM122532 (to K.M.W.).

## Supporting Information

Sequencing data are accessible through Gene Expression Omnibus (GEO) accession number GSE283716. DMS and SHAPE reactivities calculated for all the constructs are available as Supplementary files through GEO. SRA data for all the constructs (BioProject ID: PRJNA1195085) can be accessed from NCBI using the following link: https://www.ncbi.nlm.nih.gov/geo/query/acc.cgi?acc=GSE283716

