## Supplementary Figures for "Structural determinants of inverted Alu-mediated backsplicing revealed by -MaP and -JuMP"

### Supplementary Figures and Captions

**Supplementary Figure 1:** Replicate comparisons

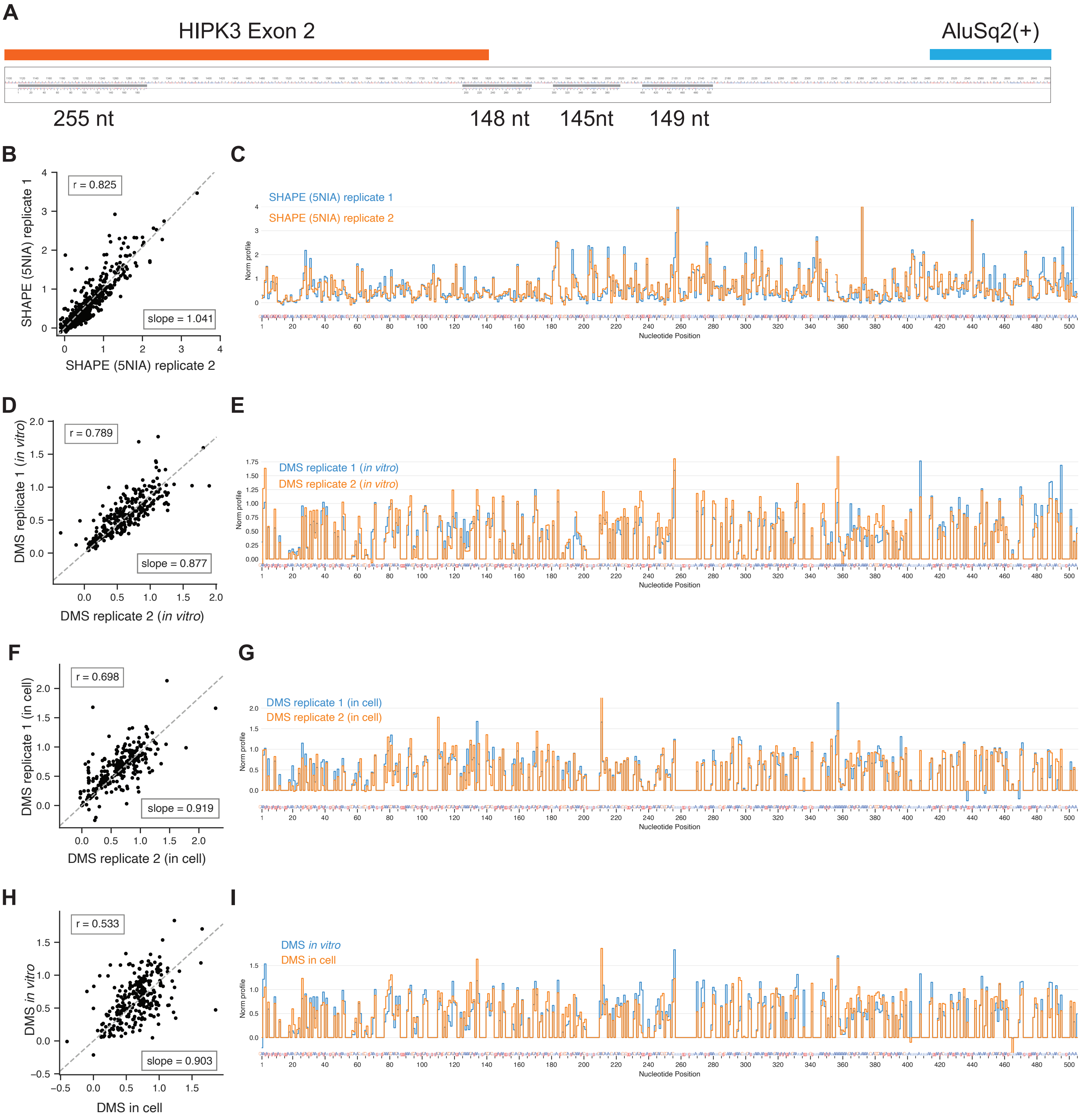

**Supplementary Figure 1:** Chemical probing replicates correlate. **A.** Gene diagram of how in cell amplicons (bottom, gray) align to the entire *in vitro* construct (top). Exon (orange) and AluSq2(+) (blue) features are colored. Aligned regions (grey, middle) are used to evaluate correlations throughout this figure to allow for comparison between all data types. **B.** Linear regression of two replicates of *in vitro* SHAPE (5NIA) probing of the wild-type construct. Values are normalized SHAPE reactivity. **C.** Skyline plot of normalized SHAPE profiles of two replicates of *in vitro* SHAPE (5NIA) data. Values are normalized SHAPE reactivity. **D.** Linear regression of two replicates of *in vitro* DMS probing of the wild-type construct. Values are normalized reactivities of only adenosine and cytidine nucleotides. **E.** Skyline plot of normalized reactivity of two replicates of *in vitro* DMS probing. Values are normalized reactivities, however guanosine and uridine nucleotides were not analyzed and set to 0 for both datasets. **F.** Linear regression of two replicates of in cell DMS probing of the wild-type construct. Values are normalized reactivities of only adenosine and cytidine nucleotides. **G.** Skyline comparison of two replicates of in cell DMS probing. Values are normalized reactivities, however guanosine and uridine nucleotides were not analyzed and set to 0 for both datasets. **H.** Linear regression comparing in cell DMS probing with *in vitro* DMS probing. Values are normalized reactivities of only adenosine and cytidine nucleotides. **I.** Skyline comparison of in cell DMS probing against *in vitro* DMS probing. Values are normalized reactivities, however guanosine and uridine nucleotides were not analyzed and set to 0 for both datasets.

**Supplementary Figure 2:** Wild-type structures

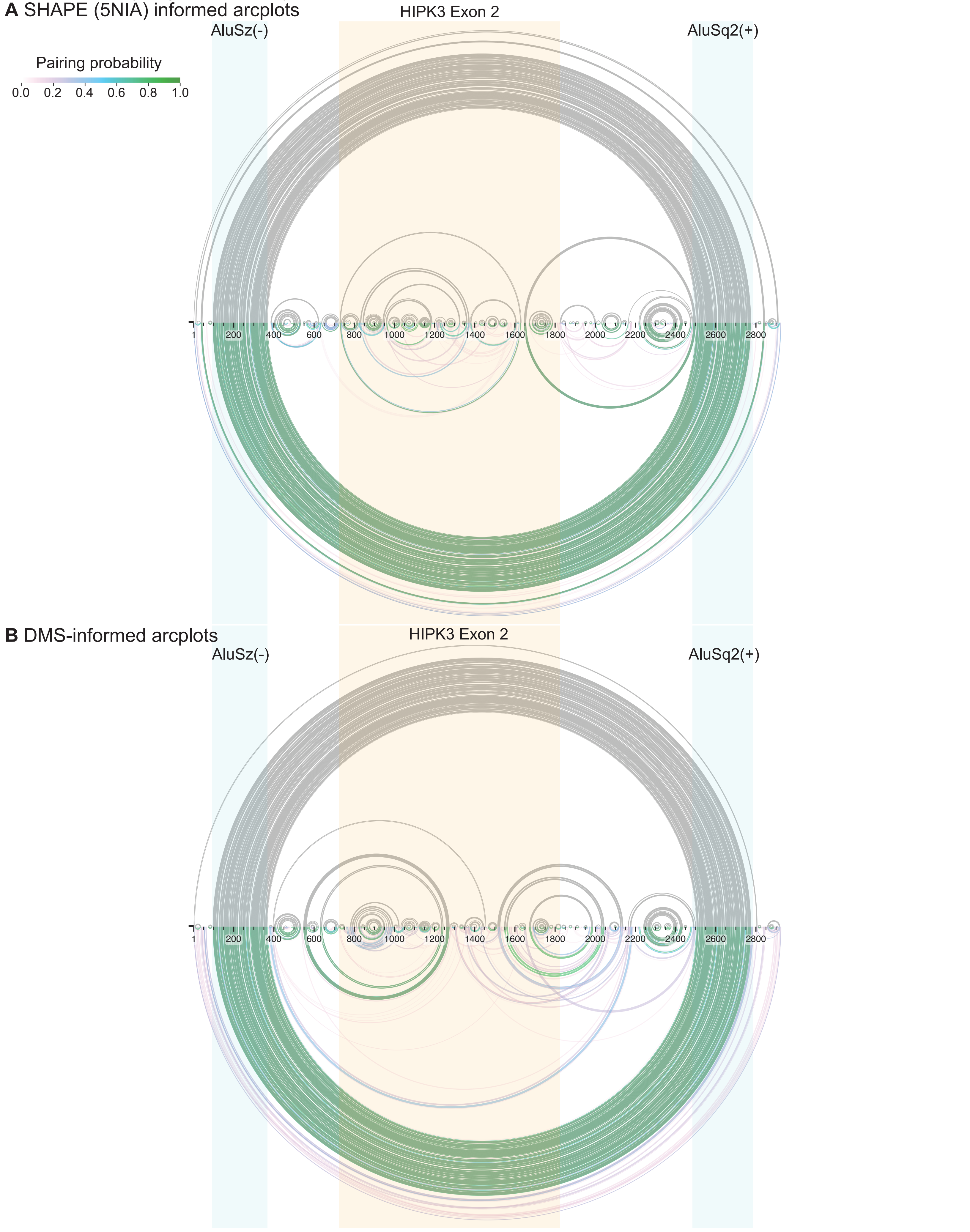

**Supplementary Figure 2:** Wild-type secondary structures (arcplots) **A.** SHAPE-informed arcplots. The blue shaded regions indicate the Alu elements, whereas the orange shaded region is the exon. The top arcs represent the minimum free energy structure, whereas the bottom represent pairing probabilities. **B.** DMS-informed arcplots. The blue shaded regions indicate the Alu elements, whereas the orange shaded region is the exon. The top arcs represent the minimum free energy structure, whereas the bottom represent pairing probabilities.

**Supplementary Figure 3:** DMS Regional Median Reactivity

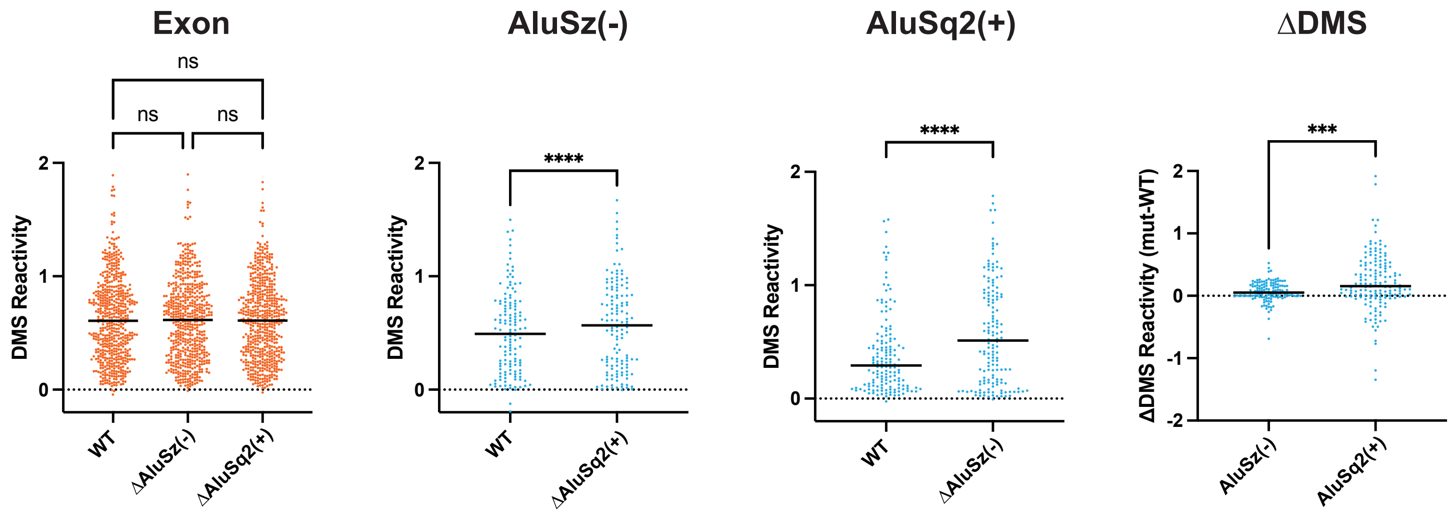

**Supplementary Figure 3:** DMS regional median reactivity. Points represent DMS reactivities of each nucleotide, with the median shown as a black line. Only adenosines and cytidines are analyzed. Significance was assessed with a one-way ANOVA for the Exon, a paired t-test for the AluSz(-) and AluSq2(+), and an unpaired t-test for ∆DMS. Significance: ns not significant, * $p < 0.05$, ** $p < 0.01$, *** $p < 0.001$, **** $p < 0.0001$

**Supplementary Figure 4:** DeltaShape

**A** AluSz(-)

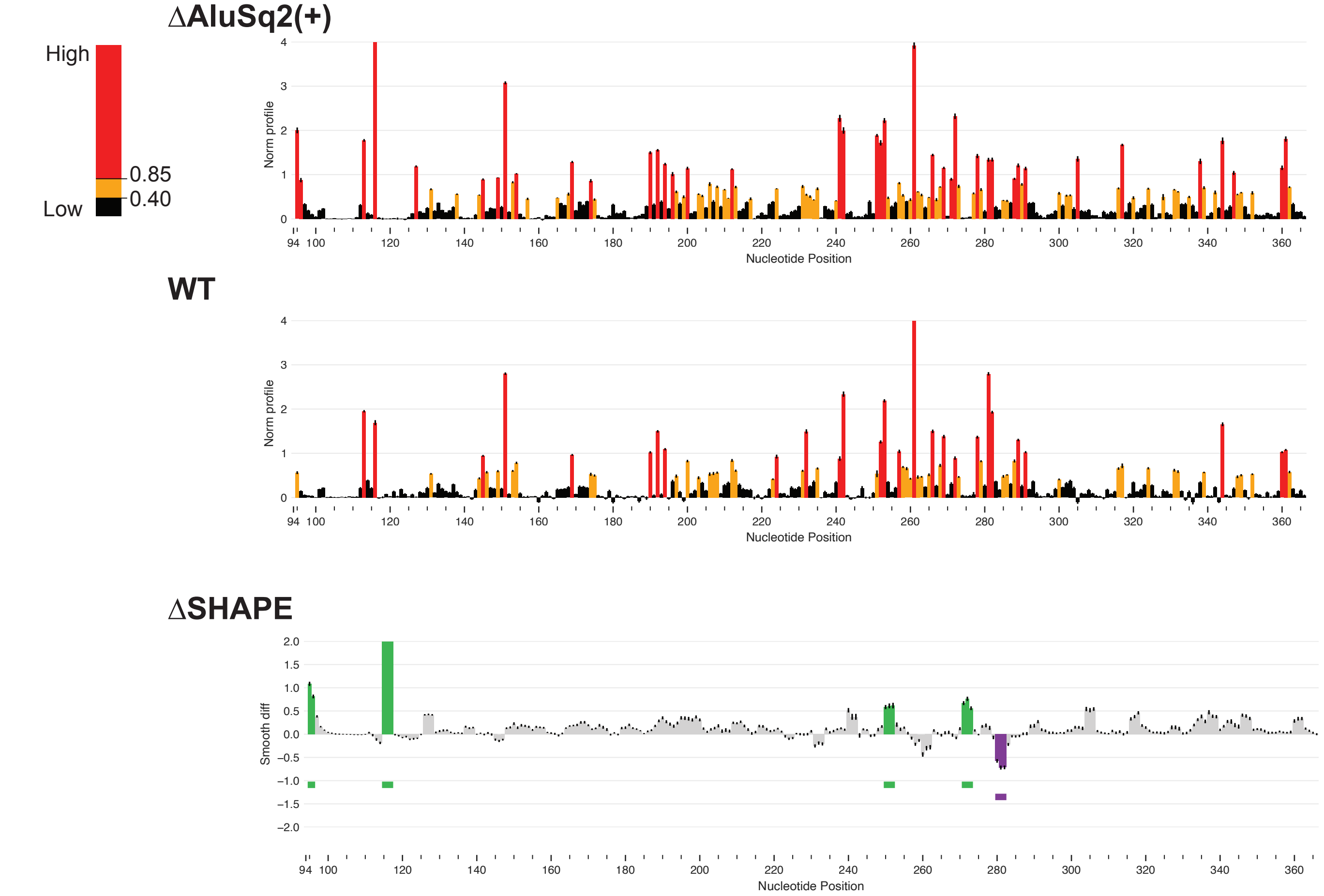


**B** AluSq2(+)

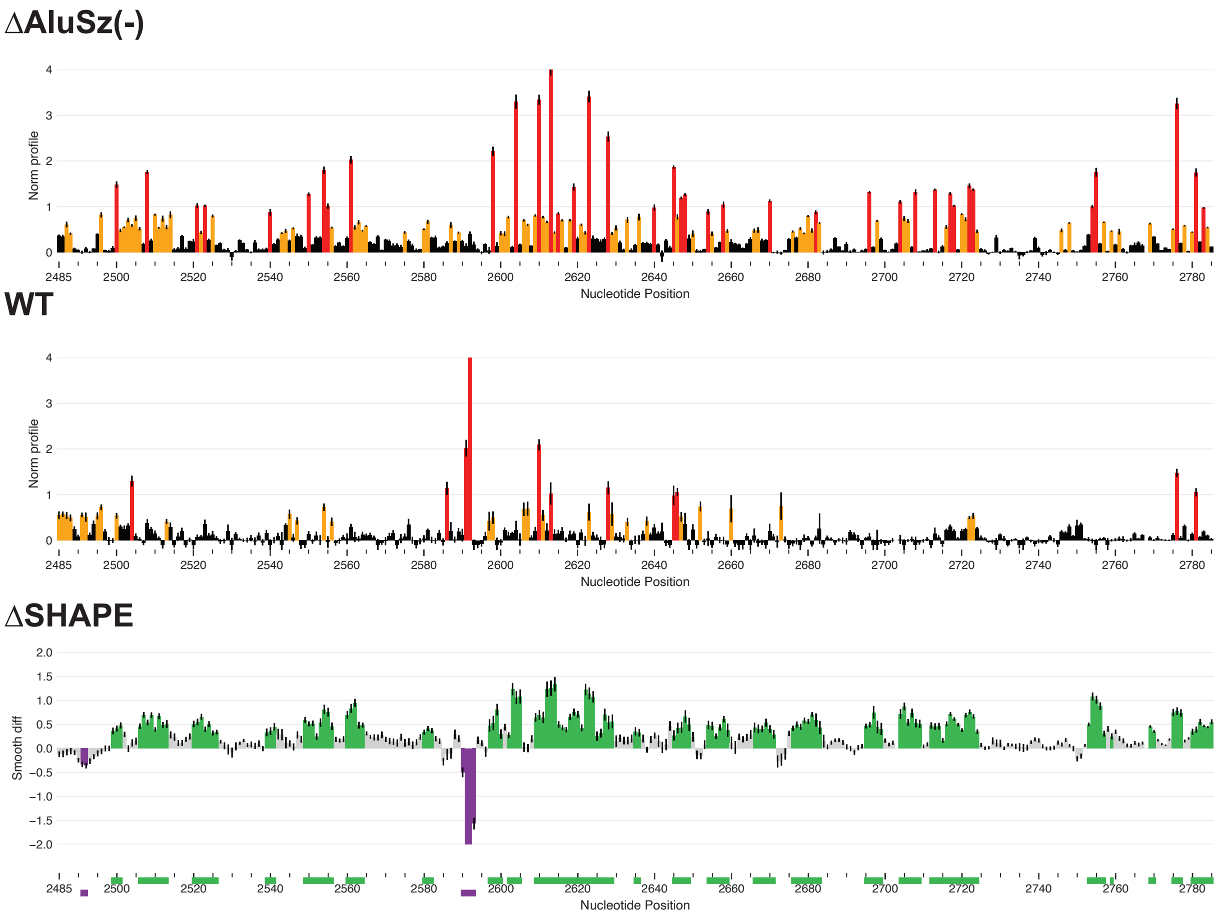

**Supplementary Figure 4:** ∆SHAPE Alu analysis **A.** ∆SHAPE analysis of AluSz(-). Top panel is the normalized SHAPE profile of AluSz(-) in the ∆AluSq2(+) context, whereas the middle panel is normalized SHAPE profile of AluSz in the wild-type context. Bottom panel is the difference between the two (∆SHAPE) across AluSz(-), (∆AluSq2(+) construct – WT construct), with statistical differences (as called by DeltaSHAPE analysis in RNAvigate) highlighted in color. Green indicates regions that have significantly increased SHAPE reactivity in the deletion construct, whereas purple indicates regions that have significantly decreased SHAPE reactivity in the deletion construct. Colored bars below the reactivity plots are highlighting the same regions. **B.** ∆SHAPE analysis of AluSq2(+). Top panel is the normalized SHAPE profile of AluSq2(+) in the ∆AluSz(-) context, whereas the middle panel is the normalized SHAPE profile of AluSq2 in the wild-type context. Bottom panel is the difference between the two (∆SHAPE) across AluSq2(+), (∆AluSz(-) construct – WT construct) , with statistical differences (as called by DeltaSHAPE analysis in RNAvigate) highlighted in color as in A.

**Supplementary Figure 5:** ΔAluSq2(+) and structure of AluSz(-) without its pairing partner

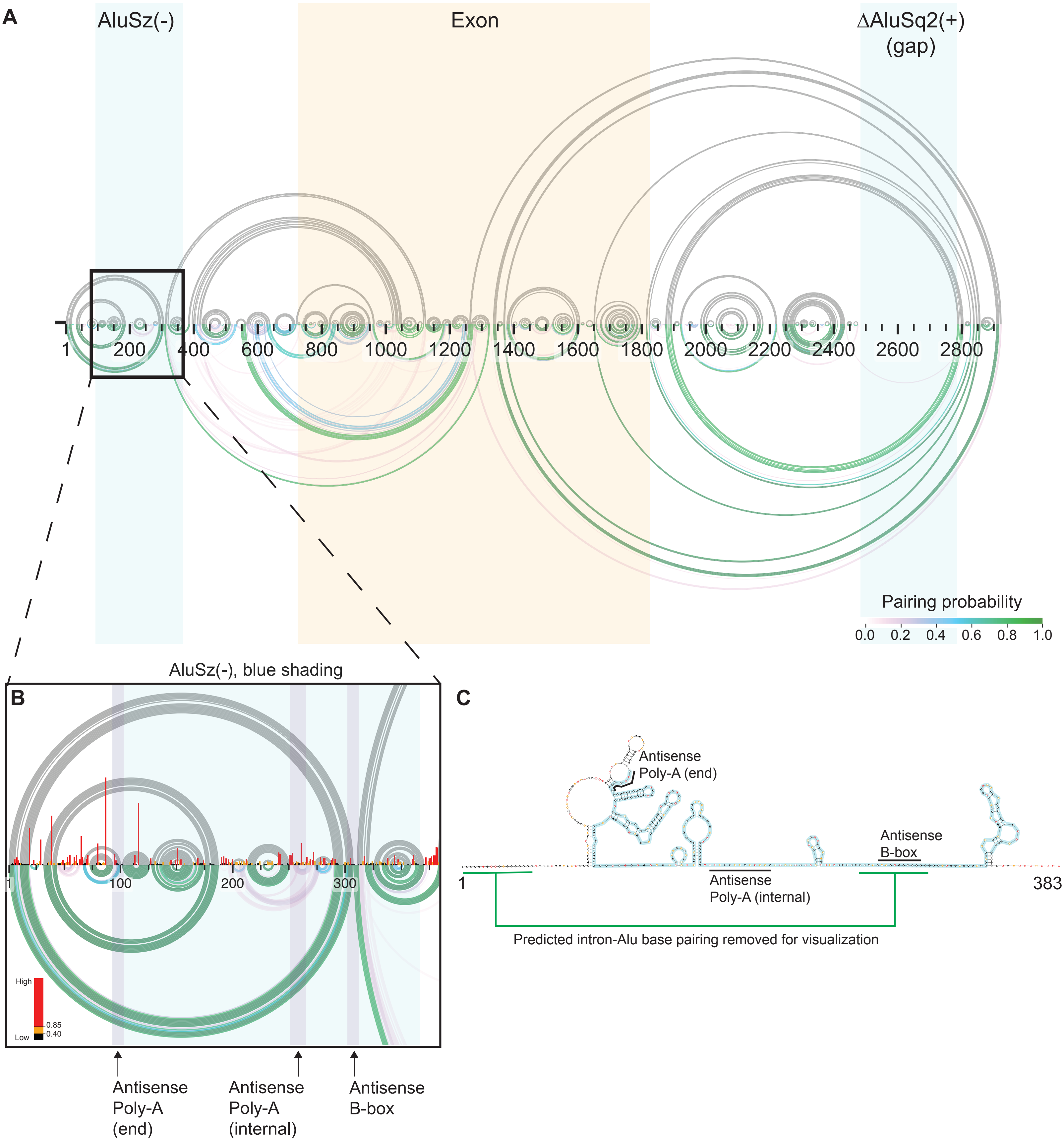

**Supplementary Figure 5:** SHAPE-informed AluSz(-) (ΔAluSq2(+)) structure **A.** SHAPE-informed arcplot structures of ΔAluSq2(+). Top (grey) arcs represent the minimum free energy structure, whereas the bottom arcs represent pairing probabilities. Blue blocks identify the Alu regions (gap indicates the deletion), whereas the orange block identifies the exon. **B.** Inset, zoomed in on the region including AluSz(-) (blue shading), with the normalized SHAPE profile. Alu element features are indicated by purple shading and annotated with arrows below. **C.** Secondary structure diagram of AluSz(-) and its flanking sequence. The Alu element is outlined in blue, and nucleotides are colored by normalized SHAPE reactivity. Alu features are annotated and denoted with a black line. Predicted intron-Alu base pairing was removed for improved visualization of the intra-Alu structure, denoted by the green line.

**Supplementary Figure 6:** ∆AluSz(-) and structure of AluSq2(+) without its pairing partner

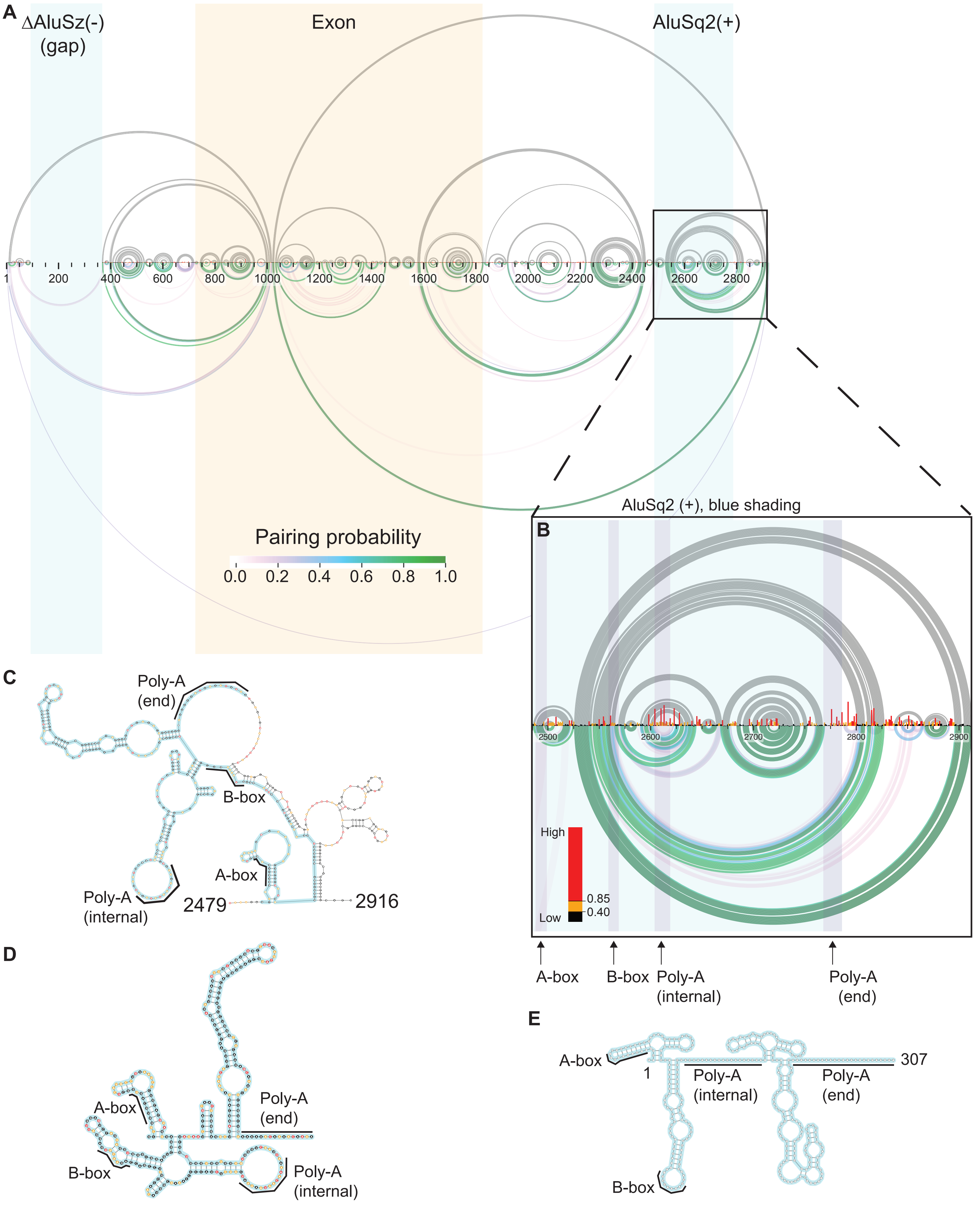

**Supplementary Figure 6:** SHAPE-informed AluSq2(+) (ΔAluSz(-)) structure **A.** SHAPE-informed arcplot structures of ΔAluSz(-). Top (grey) arcs represent the minimum free energy structure, whereas the bottom arcs represent pairing probabilities. Blue blocks identify the Alu regions (gap indicates the deletion), whereas the orange block identifies the exon. **B.** Inset, zoomed in on the region including the AluSq2(+) (blue shading), with the normalized SHAPE profile. Alu element features are indicated by purple shading and annotated with arrows below. **C.** Secondary structure of AluSq2(+) in its sequence context. The Alu element is highlighted in blue and nucleotides are colored by normalized SHAPE reactivity. Alu features are annotated and denoted with a black line. **D.** Secondary structure of AluSq2(+) folded alone, with SHAPE data but without any sequence context. The Alu element is highlighted in blue and nucleotides are colored by normalized SHAPE reactivity. Alu features are annotated and denoted with a black line. **E.** Published secondary structure of an AluSx1(+) element (31), with common Alu features annotated. The Alu element is highlighted in blue and Alu features are annotated and denoted with a black line.

**Supplementary Figure 7:** Circularization Assay schematic

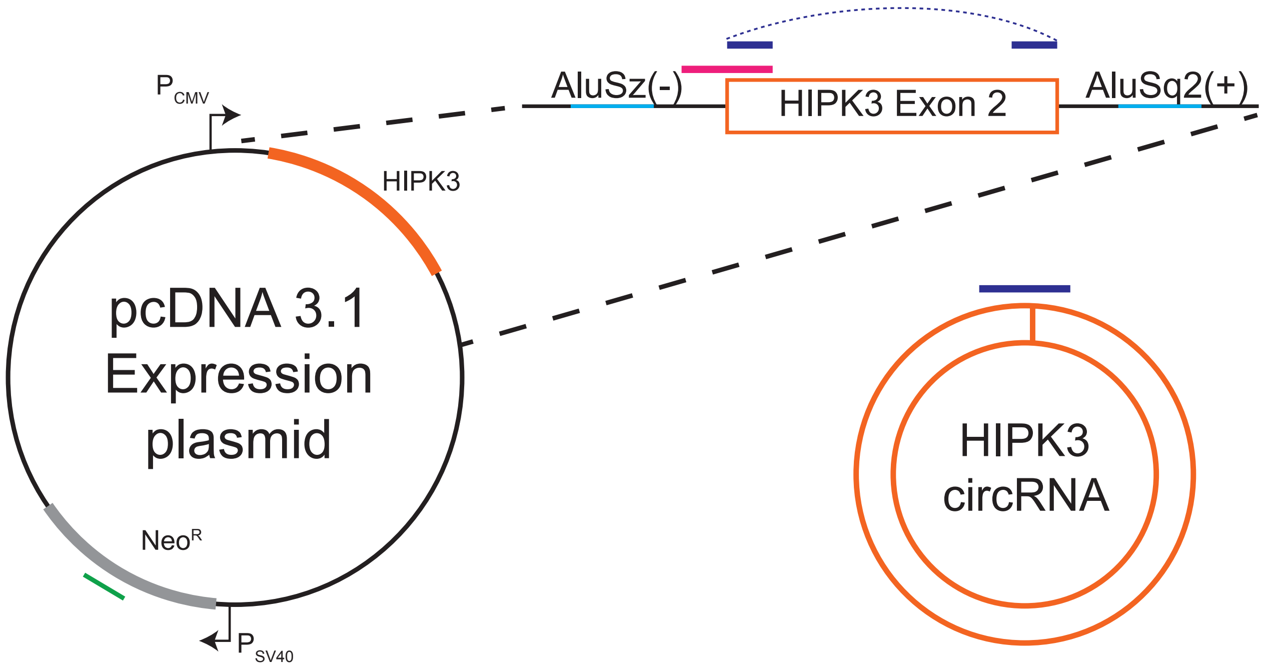

**Supplementary Figure 7:** Circularization Assay Schematic. The HIPK3 construct is expressed from a CMV promoter in the pcDNA 3.1 expression plasmid. One primer set specifically amplifies the circular form of HIPK3 exon 2 by priming outward from the exon in the linear depiction, and across the backsplice junction in the circRNA product, producing the dark blue amplicon. Another primer set specifically amplifies pre-mRNA by amplifying the intron-exon junction, producing the pink amplicon. Lastly, a control primer set amplifies the Neomycin Resistance gene on the pcDNA 3.1 plasmid, producing the green amplicon, for normalization to account for any variance in transfection efficiency.

**Supplemental Figure 8:** JuMP replicates

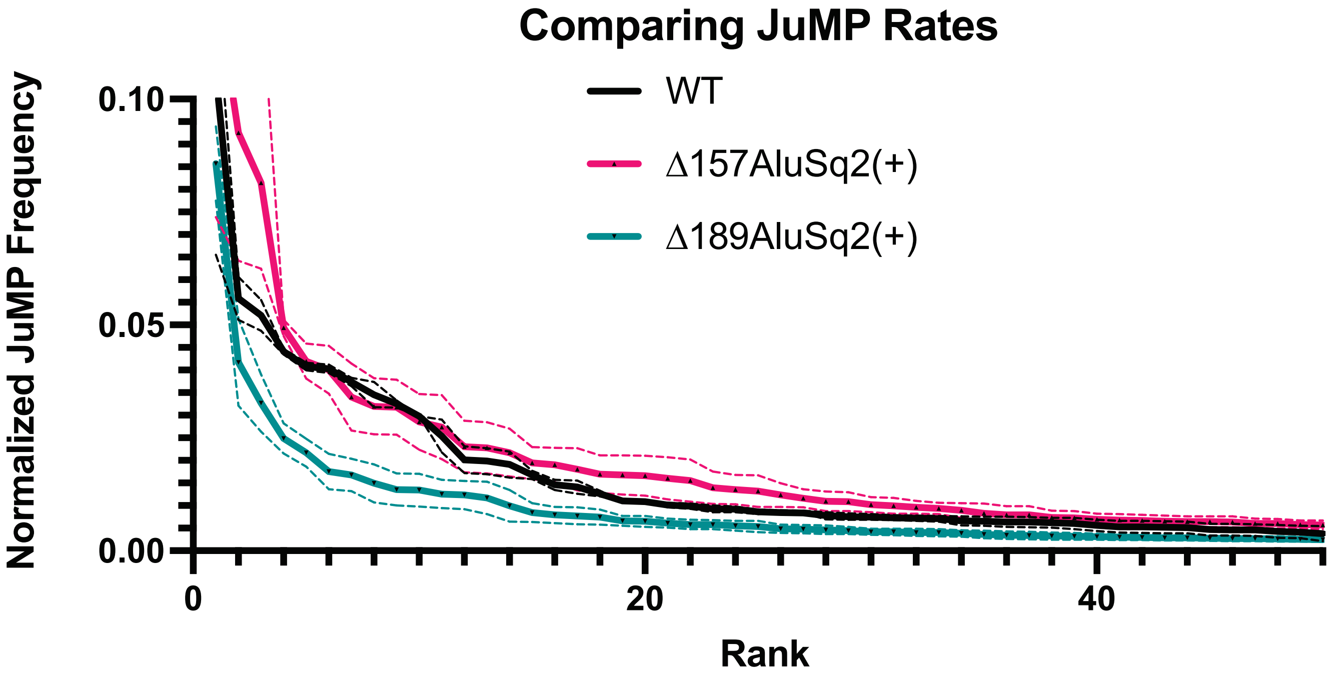

**Supplementary Figure 8:** JuMP replicate comparison. Solid lines show the normalized JuMP frequency of the highest frequency JuMP events for each construct (WT black, ∆157AluSq2(+) pink, ∆189AluSq2(+) teal). Dashed lines represent the range of two replicates. Rank denotes the order of JuMP events from the highest frequency to the lowest.
